# A Fluorogenic Array Tag for Temporally Unlimited Single Molecule Tracking

**DOI:** 10.1101/159004

**Authors:** Rajarshi P Ghosh, J Matthew Franklin, Will E. Draper, Quanming Shi, Jan T. Liphardt

## Abstract

Cellular processes take place over many timescales, prompting the development of precision measurement technologies that cover milliseconds to hours. Here we describe ArrayG, a bipartite fluorogenic system composed of a GFP-nanobody array and monomeric wtGFP binders. The free binders are initially dim but brighten 15 fold upon binding the array, suppressing background fluorescence. By balancing rates of intracellular binder production, photo-bleaching, and stochastic binder exchange on the array, we achieved temporally unlimited tracking of single molecules. Fast (20-180Hz) tracking of ArrayG tagged kinesins and integrins, for thousands of frames, revealed repeated state-switching and molecular heterogeneity. Slow (0.5 Hz) tracking of single histones for as long as 1 hour showed fractal dynamics of chromatin. We also report ArrayD, a DHFR-nanobody-array tag for dual color imaging. The arrays are aggregation resistant and combine high brightness, background suppression, fluorescence replenishment, and extended choice of fluorophores, opening new avenues for seeing and tracking single molecules in living cells.

## Introduction

Single molecule studies can reveal the operating principles of cellular machines^1, 2^ but are difficult to perform in living cells.^3, 4^ One problem is that fluorescent proteins (FPs) emit only a finite number of photons before bleaching, complicating data analysis and setting a fundamental limit on the description of biological systems. Consider a molecular machine traversing multiple microenvironments, while performing a series of operations and interacting with a succession of regulatory factors; a continuous **X**(t) track covering the entire process would give insight into reaction kinetics, the state space, transition probabilities, the noise structure, and the temporal order of molecular events.

One way to address limited observation times is to fuse multiple FPs to a target^5^ or to recruit FPs to arrays, as is done to track single RNA molecules and DNA loci.^6–10^ The RNA (or DNA) array approach can be extended to protein arrays, as demonstrated by the ‘SunTag’, an array of short epitopes that binds to single-chain GCN4 antibodies (scFv-GCN4).^11^ With such an approach, the key requirement is to maximize array occupancy while minimizing background signal by balancing relative expressions of the array and free binders. The SunTag reduces background fluorescence through spatial sequestration of free binders to the nucleus, presenting a challenge to nuclear protein tracking due to high backgrounds.^12^ The FP11-tag^13^ system reduces backgrounds using split-FPs, which have extremely slow maturation rates (~hours^14^) and whose association is essentially irreversible.^15–17^ These characteristics allow FP11 to be used for bulk visualization of native proteins^18^ and advanced imaging such as complementation activated light microscopy^16^, but prolonged single molecule tracking (SMT) has not been not demonstrated.

We developed a class of genetically encoded array tags (Figure 1a-c) based on camelid nanobodies^19, 20^ and validated their intracellular performances. We demonstrate that low background fluorescence can be achieved using a fluorogenic array, where the fluorescence of the binder FP is minimal until it binds the array. One of the nanobody arrays, ArrayG, caused a 15-fold enhancement of wild-type GFP (wtGFP) fluorescence upon binding. A fully occupied 24x-GFP-nanobody array is thus ~360 fold brighter than a single wtGFP molecule. This approach yielded high signal to noise ratio (SNR) and allowed prolonged tracking of single molecules. To efficiently track single molecules in the nucleus, we designed a nuclear-specific derivative of ArrayG, termed ArrayGN. Under some imaging conditions, the continuous exchange of binders at the array with free mwtGFP allowed temporally unlimited tracking of single molecules without decay of the overall signal from the array for thousands of frames. A third array based on DHFR nanobody (ArrayD) allowed simultaneous dual color imaging

**Figure 1.**
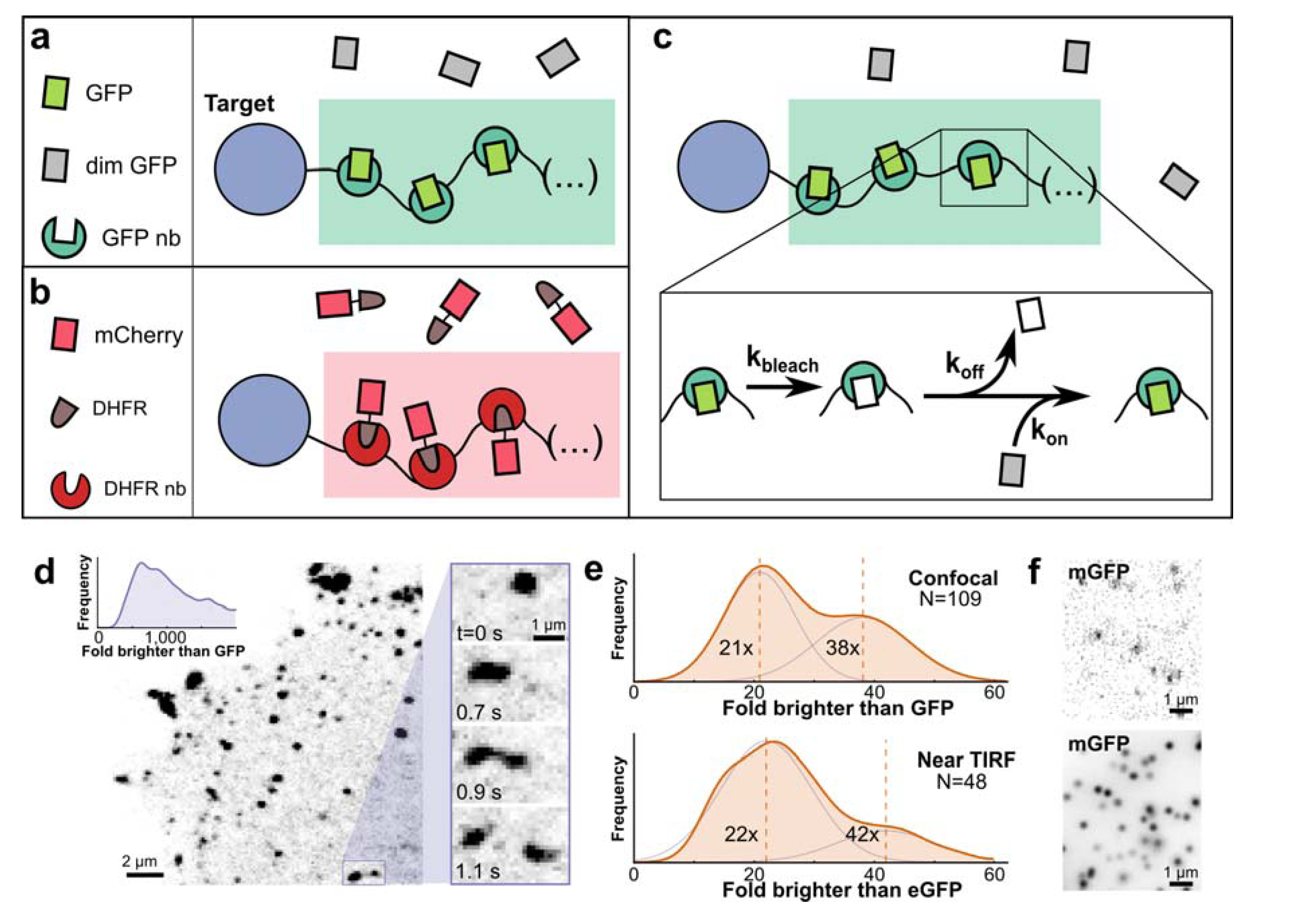
A dynamic recruitment based approach for prolonged imaging and abolishing aggregation propensity. (**a**) A schematic representation of dynamic recruitment on the the fluorogenic GFP-nanobody array. (**b**) A schematic representation of dynamic recruitment of DHFR-mCherry on DHFR-nanobody array, (**c**) Depiction of fluorescence sustenance on an array through exchange of bound photo-bleached fluorophores with dim unbound fluorophores which subsequently brighten upon binding (**d**) Representative laser-scanning confocal image of a cell co-expressing kinesin-1(KIF560)- (GBP1_24x_ GFP-nanobody array) along with eGFP as the binder. Right inset: Zoom-in showing droplet-like temporal dynamics of kinesin aggregates. Top left inset: Distribution of intensities of spots in terms of fold brighter than eGFP, showing extensive aggregation. The x-axis has been truncated at 2,000 eGFPs to emphasize the large size of even the smallest aggregates. (**e-f**) The use of monomeric A206K GFP (mGFP) abolishes aggregation. KIF560-GBP1_24x_ intensity distributions represented as fold-brighter than eGFP, show peaks corresponding to KIF560-GBP1_24x_ monomer and dimer for confocal (top) and HiLo-TIRF (bottom) (e), with representative images (**f**). Intensity distributions were fit to a two-Gaussian mixture model to estimate the local maxima for the monomeric and dimeric peaks. Vertical lines in intensity distributions represent the mean value.

The primary consideration when using these array tags is their large size, which could restrict their applicability to large molecular complexes, or machines (e.g. kinesins) that have evolved to drag large payloads. However, both the moderately sized integrin β1 (~88 kDa) and histone H2B (~14 kDa) tolerated the ArrayG/GN tag in a multitude of functional assays. For instance, β1-ArrayG restored contractility to an integrin knockout cell line. The fluorogenic nature of the tag will potentially allow the use of shorter repeat arrays, where a tradeoff between array length and retention of native functionality will define the maximum achievable track duration.

## Results and Discussion

### Preliminary scan of possible array / binder pairs

We identified six high affinity protein interaction pairs in the literature (**Supplementary Table 1**). Five were nanobody-antigen pairs. The small size and monomeric nature of nanobodies facilitates generating tandem repeat arrays.^19^ The intracellular recruitment potential of each interaction pair was quantified using a nuclear compartmentalization assay, which was needed since *in vivo* affinities in cellular compartments can differ significantly from values determined *in vitro* (**Supplementary Note 1, Supplementary Figure S1, Supplementary Table 1**). We chose to further develop GBP1-GFP and NB113-DHFR, the two best performing interaction pairs. In particular, the GBP1-GFP binder pair provided a potentially attractive candidate for background fluorescence suppression due to GBP1’s ability to enhance^21^ wtGFP fluorescence upon binding.

### Barrier 1: Aggregation

All multivalent recruitment strategies must contend with the possibility of aggregation. We encountered aggregation in our preliminary tests of the GBP1 nanobody array. In these early tests, we had fused a 24x repeat of GPB1 (GBP1_24x_) to KIF560, a (+) end directed kinesin-1 protein lacking the cargo-binding domain.^5, 22, 23^ Co-expression with eGFP resulted in numerous bright spots exhibiting complex dynamic behavior (**Supplementary Movie S1**.However, spot intensity analysis using calibrated confocal microscopy revealed higher-order aggregates of KIF560-GBB1_24x_, with a typical spot containing ~1000 eGFPs (Figure 1d, **Supplementary Figure S2**).

### Addressing barrier 1: monomeric GFP

Aggregation was resolved by replacing eGFP, which has a weak dimerization propensity (K_d_ = 110 μM)^24^, with the monomeric A206K^24^ mutant (‘mGFP’). This modification collapsed the previously broad distribution of intensities into two peaks with maxima corresponding to 21 and 38 GFPs (confocal microscopy) or 22 and 42 GFPs (HiLo-TIRFM) (Figure 1e-f). This likely represents a mixture of GBP1_24x_-KIF560 homodimers and heterodimers of native kinesin 1 and GBP1_24x_-KIF560.

### Barrier 2: Low throughput and variable background signal

The two requirements for attaining high signal-to-noise ratio (SNR) using a dynamic array strategy are high array occupancy and low background fluorescence. Although GBP1_24x_/mGFP allowed single molecule tracking in living cells, experimental throughput was initially low. Some cells were useable and provided good tracking data, but the vast majority of cells either had high backgrounds or poor signal.

### Addressing barrier 2: Binding induced fluorescence enhancement for background suppression

To reconcile low background and high array occupancy, we explored a fluorogenic system in which where the free fluorophore is dark and only becomes fluorescent upon binding to the array. Single GBP1 nanobodies have been previously shown to enhance the fluorescence of wtGFP upon binding.^21^ We hypothesized that replacing mGFP with a monomeric A206K wtGFP (mwtGFP) would reduce background from free binders.

To compare mwtGFP and mGFP fluorescence, we expressed either mwtGFP or mGFP fused to mCherry via a P2A^25^ self-cleavable peptide (Figure 2a). Flow-cytometric analysis of the mCherry-normalized basal fluorescence showed ~30 fold less fluorescence for free mwtGFP compared to mGFP (**Supplementary Figure S3**). In contrast to FACS, HiLo-TIRFM imaging revealed an 18-fold weaker fluorescence for mwtGFP (5.5% of the brightness of mGFP) (Figure 2b).

**Figure 2.**
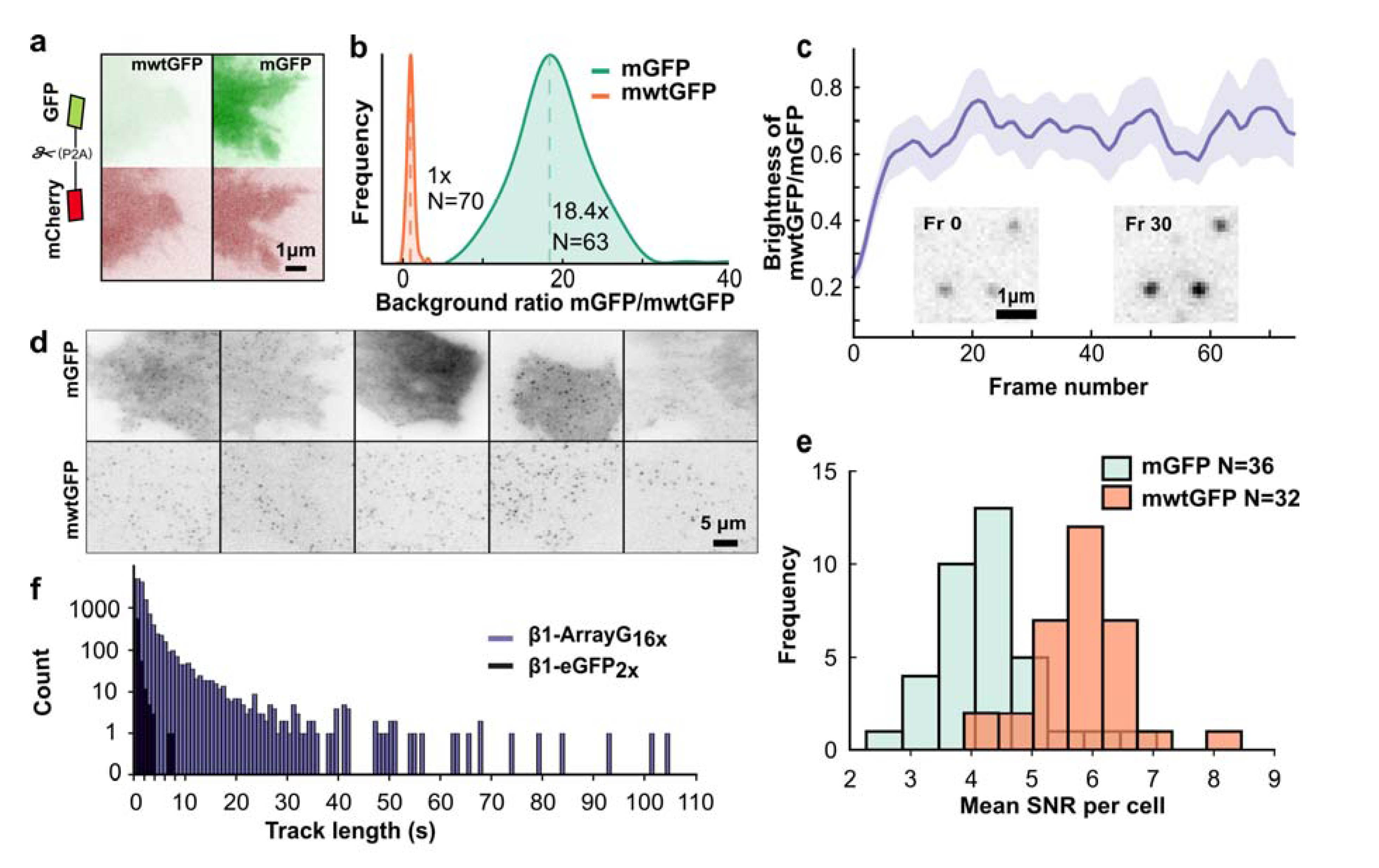
Fluorescence enhancement of mwtGFP upon binding GBP1 array suppresses background fluorescence. (**a**) TIRF images of HeLa cell lines stably expressing self-cleavable constructs for ratiometric normalization of GFP fluorescence: mCherry-p2a-mwtGFP (left) or mCherry-p2a-mGFP (right). (**b**) Distribution of mGFP intensity (N=63 cells) normalized by mean mwtGFP intensity (N=70 cells) measured using TIRF. Vertical dashed lines represent mean value. (**c**) Time dependence of GBP1_2_4x array bound mwtGFP brightness relative to mGFP. Inset: TIRF images of mwtGFP+GBP1_24x_ spots at frame 0 and frame 30 (at 20Hz image frequency), showing significant brightening. (**d**) TIRF images of U2OS cells stably expressing integrin β1-ArrayG_16_x and either mGFP (top) or mwtGFP (bottom) binder. (**e**) Histogram of the mean array signal-to-noise per cell for integrin 1-ArrayG_16_x co-expressing either mGFP or mwtGFP. (**f**) Trajectory length distribution for β1-eGFP2x (659 trajectories from 4 cells) and β1-ArrayG_16x_ (13,423 trajectories from 4 cells). Minimum trajectory length is 10 frames.

To measure the magnitude of the fluorogenic effect in the context of the GBP1 array, we co-expressed KIF560-GBP1_24x_ arrays in cells with either monomeric A206K wtGFP (mwtGFP) or mGFP. In fixed cells GBP1_24x_ was ~75% as bright when occupied with mwtGFP than when occupied with mGFP (Figure 2c). Interestingly mwtGFP-bound GBP1_24x_ brightened upon illumination even at relatively low laser powers (< 0.01 kW cm^2^) and photo-matured completely within ~200-300 ms (Figure 2c, inset). From these measurements we conclude that individual mwtGFPs undergo a net ~15-fold fluorescence enhancement upon binding the array (relative to mGFP, mwtGFP is 75% as bright when bound to GBP1 and 5.5% as bright when unbound).

To test whether the SNR of localized arrays was significantly improved by using mwtGFP, we compared cell lines expressing integrin β1-GBP1_16x_ and either mwtGFP or mGFP. We chose integrin β1 over Kif560 to spatially confine the GBP1 scaffold to a single cellular compartment, facilitating quantitative comparison in live cells. Nearly every cell expressing mwtGFP exhibited arrays with high SNR (**Supplementary Movie S2**), whereas for mGFP single molecule detection was obscured in a large fraction of cells due to high background fluorescence (Figure 2d). Even when comparing mwtGFP cells to the subset of mGFP cells where single molecules could be detected, SNR increased significantly from 4.3 for mGFP to 5.8 for mwtGFP (Figure 2e). Moreover, when tracked under identical conditions, β1-GBP1_16X_ markedly increased track duration compared to β1-eGFP_2X_ (Figure 2f) allowing tracking up to 105 seconds (2100 frames, **Supplementary Movie S3**). For ease of discussion, we refer to the GBP1_Nx_ + mwtGFP as ‘ArrayG_Nx_’, where N represents the number of tandem repeats in the array.

### Barrier 3: Photobleaching imposes a limit on trajectory length

Despite the extremely long track lengths of ArrayG, the tracks were still finite. However, a simple theoretical perspective on fluorescent binders that are stochastically exchanging on an array implies that in certain kinetic regimes temporally unlimited tracking is possible. Specifically, the average number of unbleached fluorophores *F* bound to an array with *N* binding sites will depend on the effective on rate k_on_, the off rate k_off_, and the bleach rate k_b_ (see **Supplementary Note 2** for derivation).

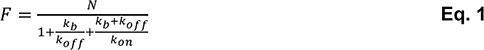

Using Eq. 1, we plotted the steady state array occupancy as a function of k_off_ and k_bleac_h assuming a large constant k_on_. The plot shows a large parameter space in which *F* can be driven close to *N,* given k_off_>>k_bleach_ and a non-limiting *kon* (Figure 3a). Experimentally, the photo-bleaching rate is primarily determined by exposure duration and frequency. By modulating exposure frequency one should be able to control bleach rate such that array fluorescence is indefinitely sustained.

**Figure 3.**
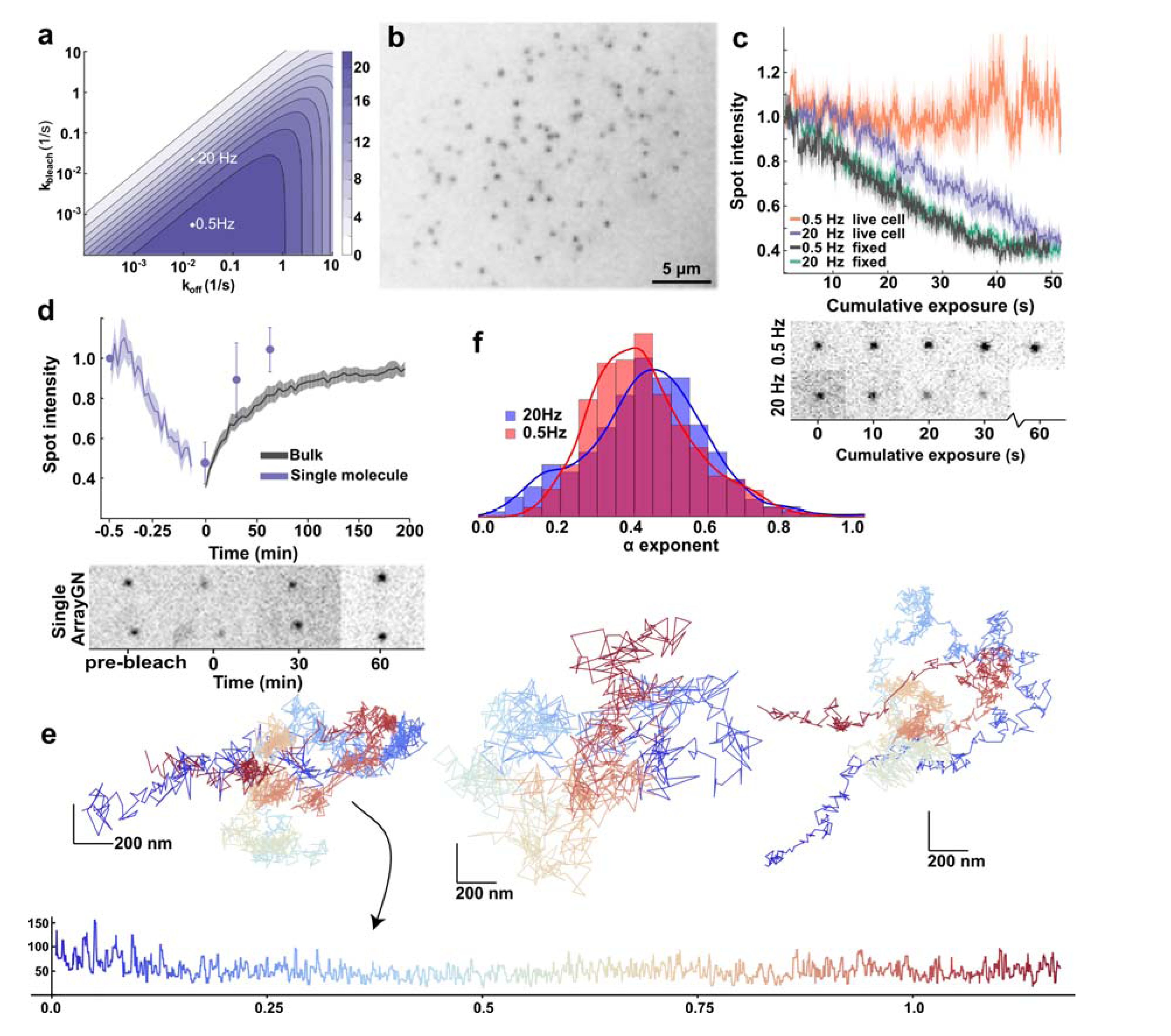
Hour-duration tracking of single histones in the nucleus using ArrayGN. (**a**)Theoretical prediction of array occupancy by unbleached fluorophores as a function of bleach rate, k_bleach_, and unbinding rate, k_off_ (plotted using k_on_=10). The occupancy estimations for 20 Hz and 0.5 Hz imaging are shown as white crosses. (**b**) HiLo-TIRF image of a nucleus with integrated H2B-ArrayGN_24x_. (**c**) *Top:* Normalized spot intensity for tracked single-molecules as a function of cumulative laser exposure for 20 Hz or 0.5 Hz (50 ms exposure/frame) in fixed and live cells. (0.5 Hz live: N=20; 0.5 Hz fixed: N=24, 20 Hz live: N=19; 20 Hz fixed: N=23). Solid lines are mean intensity, shading is s.e.m. *Bottom:* raw images of single spots of H2B-ArrayGN_24x_ for 20 Hz and 0.5 Hz in live cells. (**d**) *Top*: FRAP for single H2B-ArrayGN_24x_ tracked over time (N=3 traces) and bulk FRAP from nuclear regions in cells overexpressing H2B-ArrayG_N_ and wtmGFP (N=5 cells, 2 ROIs per cell). Solid line is mean intensity, shading and error bars are s.e.m. *Bottom:* raw image of single H2B-ArrayGNN_24x_ FRAP, (**e**) Three examples of single histone trajectories, color coded by time. Bottom: A five-point median-filtered plot of a single histone’s displacements over time. (**f**) Distribution of α values for H2B-ArrayGNN_24x_ imaged at 0.5Hz and 20 Hz.

### Addressing barrier 3:Temporally unlimited trackability

Chromatin embedded fiducials are relatively immobile (compared to processive molecular motors and integrins), facilitating the evaluation of unlimited trackability. We chose histone H2B due to its extremely slow dissociation kinetics (multiple hours).^26^ For efficient nuclear import and integration of an H2B-ArrayG fusion, we generated ‘ArrayGN’, a modified version of ArrayG in which 3 copies of Snurportin-Importin (3 binding domain (IBB)^27^ were interspersed throughout the array (H2B-(Glycine-Serine repeat)_20x_-(GBP1_8x_-IBB)_3x_) (Figure 3b, **Supplementary Movie 4**).

We compared the photo-stability of H2B-ArrayGN in G1/S synchronized cells (**Methods**) imaged under continuous laser exposure versus pulsed exposure (50 ms exposure at 0.5 Hz) (Figure 3b, c, **Supplementary Movie 5**) using identical laser power settings. Continuous illumination resulted in progressive decay of single H2B-ArrayGN intensities, whereas imaging under the pulsed exposure format showed no signs of intensity decay (Figure 3c, **Supplementary Movie 5**). Furthermore, when imaged after formaldehyde fixation of cells, spot intensity decayed similarly for both continuous and pulsed exposure (Figure 3c), demonstrating that stochastic turnover on the array was the fundamental mechanism of signal replenishment. Finally, we directly observed fluorescence recovery after photo-bleaching in both ensemble (t_1/2_ = 27±9 min) and single-molecule modes in cells stably expressing H2B-ArrayGN24X+ mwtGFP (t_1/2_= 23±4 min) (Figure 3d, **Supplementary Figure S4, Methods**), demonstrating that a fully bleached array will gradually recover its fluorescence due to stochastic turnover.

### Intracellular performance tests and tag validation

Having developed a tag (ArrayG_24x_) 360 fold brighter than wtGFP with three defining features, (1) aggregation resistance, (2) background suppression through fluorogenicity, and (3) indefinite trackability, we tested its performance in living cells.

### Test 1: Multiscale dynamics of single chromatin bound histones in the living nucleus

We measured the dynamics of single H2B-ArrayGN24x labeled nucleosomes in G1/S synchronized cells. The ArrayGN_24x_ tags with their low backgrounds allowed sustained tracking of H2Bs and yielded a large SMT dataset. For example, one cell with 194 H2B-ArrayGN_24X_ molecules provided 147,000 seconds of precision 2D tracking data (0.5 Hz imaging, mean track length ~760 seconds, maximum track length 4200 seconds). At the lower imaging rate (0.5Hz), track lengths were no longer limited by loss of signal due to photobleaching; a track typically ended when a molecule moved out of the HiLo imaging volume. XY(t) data for three histones are shown in Figure 3e. The single histone trajectories had rich dynamics; for example, the nanometer vs. hour plot (bottom of Figure 3e) shows the displacements of one histone’s position over more than 1 hour, encompassing a 4x range of frame-to-frame displacements (Figure 3e, bottom).

While the technology should facilitate the detailed study of specific gene loci and chromatin dynamics (such as looping and TADs formation), our focus here are the baseline characteristics of living chromatin, especially its spatial and temporal geometry. Sequencing methods such as Hi-C suggest that chromatin is spatially self-similar at length scales from ~100 kilobases to several megabases.^28, 29^ Separately, there is evidence that chromatin is also temporally fractal; optical tracking of single loci in prokaryotes and eukaryotes suggests a Hurst parameter *H* ~ 0.25 (corresponding to an exponent α = *2H* ~ 0.5).^30–33^ H2B-ArrayGN imaged at 20 Hz and 0.5 Hz were fit to fractional Brownian motion (fBm) using a maximum likelihood criterion. The 0.5 Hz tracks yielded an exponent α = 0.42 ± 0.14 (95% confidence interval [0.41, 0.43]) while the 20 Hz tracks yielded α = 0.43 ± 0.16 (95% confidence interval [0.42, 0.45], **Figure 3f, Methods**). Peripheral histones (nearer the nuclear envelope) were more confined than ‘interior’ histones (**Supplementary Figure S5**). The α and apparent diffusion coefficient estimates obtained from the chromatin-integrated ArrayGN-histones are in excellent agreement with the values obtained from short tracks of single GFP-tagged histones by Shinkai and coworkers.^34^

The similarity of the α values for different systems (single GFP-H2B vs. H2B-ArrayGN at 20 Hz and 0.5 Hz) suggests that the labeled histones are reporting the underlying motions of relatively large regions of the chromatin, whose molecular weight far exceeds that of any single protein (or protein complex). It is notable that recent estimates of α in eukaryotes tend to be closer to 0.4 than 0.5, but these differences are well within the variation of ± 0.1 seen across cell lines.^35^ The fact that the short (20 Hz) trajectories, covering seconds to minutes, and the hour-timescale (0.5Hz) trajectories give nearly identical α values strongly supports the notion that chromatin has a fractal temporal structure.

### Test 2: Molecular motors/Kinesins

A second major application area for live cell SMT is the large field of motor mechano-chemistry and regulation. In contrast to the tethered nature of H2B dynamics, motor proteins exhibit relatively fast directional motion. To inspect the intricate details underlying the fast, directionally-persistent motion of motor proteins, we tracked KIF560-ArrayG24x using HiLo-TIRFM at 20 Hz. This yielded hundreds of directionally processive trajectories per cell, spanning up to several μm in length (Figure 4a, **Supplementary Movie S6**). Live staining of cells with the fluorogenic SiR-tubulin dye allowed us to track single molecules of kinesin on microtubule track for extended lengths of time at higher imaging rates (80Hz) (Figure 4b, **Supplementary Movie S7**). The speed of KIF560-ArrayG_24x_ motors was sharply distributed about a mean of 1.48 μm/s (Figure 4c-d). The average speeds and run-lengths also matched existing live cell measurements.^5, 11^

**Figure 4.**
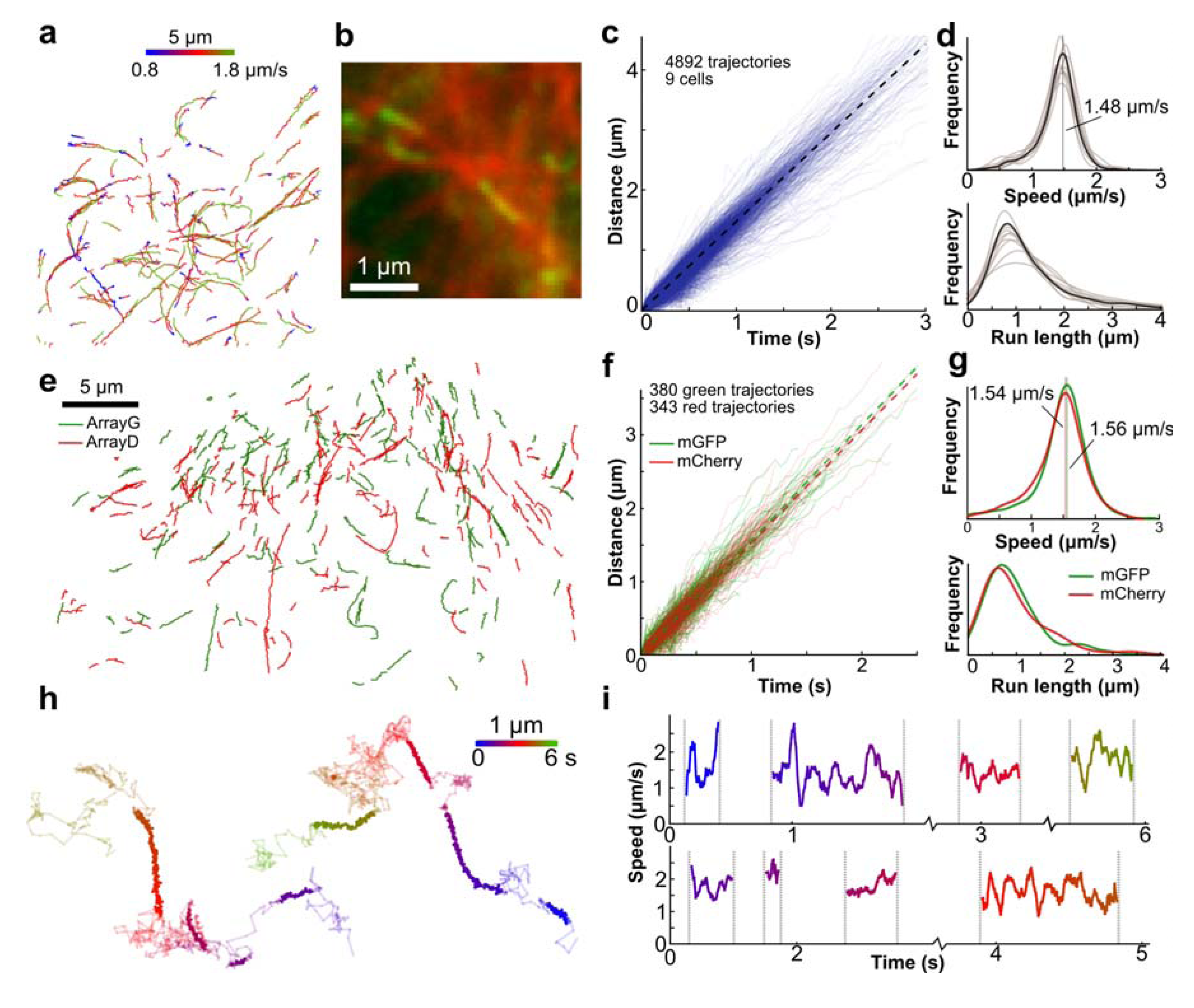
Kinesin-ArrayG trajectories demonstrates proper functionality and reveals large speed fluctuations. (**a**) Trajectories of single KIF560-ArrayG_24x_ motors labeled with mwtGFP colored by speed. (**b**) Time projection of KIF560-ArrayG24x at 80 Hz on labeled tubulin. (**c**) Distance versus time trace for 4892 individual trajectories from 9 cells. Dashed line represents the mean speed. (**d**) Distribution of trajectory speeds (top) and run lengths (bottom) from c. Individual cell distributions are shown in transparent, with the mean as solid. Vertical dashed line indicates the mean (1.5 μm/s). (**e**) Trajectories of kinesin in cells co-expressing KIF560-ArrayG24x + mwtGFP (green) and KIF560-ArrayD24x + DHFR-mCherry (red). (**f**) Distance versus time trace of trajectories shown in e. (**g**) Distribution of speed and run length from f. (**h**) A high frame-rate trajectory of KIF560-ArrayG (180Hz) colored by time. Linearly persistent trajectory segments are colored bold. (**i**) Instantaneous speed of the linearly persistent segments from the trajectories shown in h.

To extend the scope of array-based tags for simultaneous multi-color imaging, we developed an orthogonal array tag based on the nb113 DHFR-nanobody^36^+*E*. *Coli* DHFR array-binder pair, termed ArrayD. One added advantage of ArrayD is that it can be used to recruit any FP as a fusion to DHFR. Despite a lower reported affinity^36^ of the nb113+DHFR pair compared to the GBP1+GFP pair, ArrayD_24x_ allowed efficient tracking of single kinesins when coexpressed with DHFR-mGFP (**Supplementary Movie S8**) or DHFR-mCherry (**Supplementary Movie S9**). The corresponding average speeds and track lengths were similar (1.51 μm/s and 1.61 μm/s average speed, 0.94 μm and 1.10 μm average run length, for mGFP-DHFR and mCherry-DHFR, respectively) (**Supplementary Figure S6**). Furthermore, by introducing ArrayG_24x_ (with mWTGFP) and ArrayD_24x_ (with DHFR-mCherry) in the same cell, we were able to simultaneously track two populations of KIF560 (**Supplementary Movie S10**). These gave indistinguishable average speeds (1.54 μm/s and 1.56 μm/s for mGFP and mCherry, respectively) showing internally consistent results and demonstrating that these arrays are suitable for simultaneous multicolor imaging (Figure 4e-g).

### Highly resolved temporal dynamics of kinesin

The photo-stability and brightness of ArrayG also made it possible to capture high-frequency features of kinesin dynamics. Using a 25-fold higher laser illumination (0.2 kW/cm^2^), we were able to track KIF560 at 180 Hz, revealing up to 4 cycles of binding, directed motion, unbinding, and free diffusion in single trajectories (Figure 4h, **Supplementary Movie S11**). During the periods of directed motion, the speed varied from less than 1 μm/s to nearly 3 μm/s, with an average speed around 1.5 μm/s (Figure 4i). To our knowledge, such short-lived speed transients have not been previously reported using genetically-encoded tags. Precision, high-frequency tracking using ArrayG may help answer questions regarding adaptive changes in behavior of kinesin moving on a dynamic microtubule landscape^37^, particularly in living systems only accessible through the use of genetically encoded tags.

### Test 3: Long-term imaging of integrin β1 using ArrayG reveals novel mobility states and context-dependent state transition-rates

A third major application area of single molecule tracking is the study of transmembrane complexes. Previous single molecule studies have shown that transmembrane proteins exhibit multiple states of motion, corresponding to different functional states.^38–40^ A common limitation in these studies is trajectory duration, preventing a proper/thorough quantification of transient state-switching dynamics. To explore the utility of ArrayG for tracking transmembrane proteins, we fused it to integrin β1. Previous short time-scale SPT-PALM experiments have suggested that integrins transition between diffusive inactive conformations and immobile active conformations.^40^ We used four separate criteria to evaluate the functionality of β1-ArrayG: enrichment at focal adhesions, functional complementation in integrin β1-deficient cell lines, rate of diffusion, and response to Mn^2^+. We generated stable cell lines expressing a more compact derivative of ArrayG with 16 repeats (ArrayG_16x_) fused to integrin β1 (**Supplementary Note 3**). Through trans-complementation of cells lacking native integrin β1 with β1-ArrayG_16x_ or β1-P2A-eGFP fusions we verified that β1-ArrayG_16x_ enriched at focal adhesions (Figure 5a) and restored wild type levels of cell/substrate engagement and spreading to a spreading-deficient integrin knockout line (Figure 5b-c, **Supplementary Figure S7,Supplementary Note 3**).

**Figure 5.**
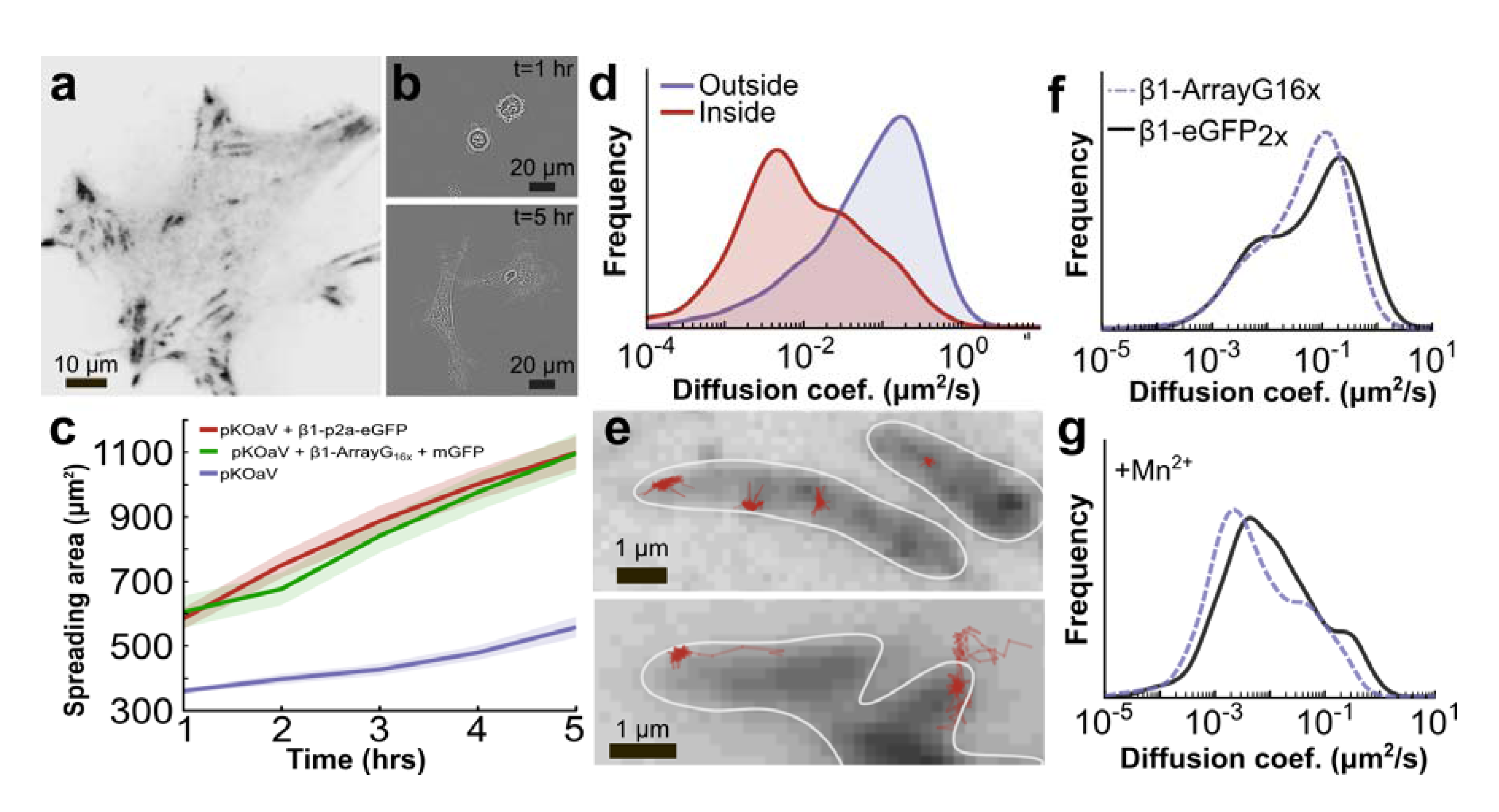
Evaluation of integrin p1-ArrayG_16x_ functionality. (**a**) Representative TIRF image of pKOαV MEF cells after induction of integrin β1-ArrayG_16x_ and mGFP. (**b**) Brightfield images of pKOαV MEFs overexpressing β1-ArrayG_16x_ + mGFP at 1 and 5 hours post seeding on fibronectin. (**c**) Mean spreading area versus time for *pKOαV* (53 cells), *pKOαV+β1-(p2a)-eGFP*(59 cells), and *pKOαV+β1-ArrayGi_6_x^+^mGFP* (57 cells). Shaded area represents s.e.m. (**d**) Diffusion coefficient distribution of integrin β1-ArrayG_16x_ inside and outside focal adhesions. N=5 cells, 1852 trajectory segments. (**e**) Examples of immobilized trajectories inside focal adhesions (top) and trajectories transitioning between diffusive and immobilized states at focal adhesion boundaries (bottom). Focal adhesion boundaries highlighted in white. (**f**) Distribution of diffusion coefficients of β1-eGFP2x *(N=4* cells, 659 trajectories) and β1-ArrayG16x (N=9 cells, 2860 trajectory segments) in untreated cells. (**g**) Distribution of diffusion coefficients after 2mM Mn^2^+ treatment (β1-eGFP_2)_: N=3 cells, 350 trajectories, β1-ArrayG_16x_: N=8 cells, 1388 trajectory segments).

Quantification of diffusivity revealed enhanced immobilization of β1-ArrayG_16x_ molecules inside adhesions marked by vinculin-mCherry (Figure 5d-e, **Supplementary Figure S8**). We observed the lateral diffusion of β1-ArrayG_16x_ across adhesion boundaries and subsequent immobilization inside the adhesion (Figure 5e, **Supplementary Movie S12**), supporting the proposed ‘archipelago’ model of focal adhesion architecture.^41^

To assess the impact of the ArrayG tag on lateral diffusion within the membrane, we compared the dynamics of (31-ArrayG_16x_ to β1-eGFP_2x_ in pKO-αVβ1cells^42^ that express endogenous integrin β1. The diffusion coefficients had a bimodal distribution representing a freely diffusing and an immobilized population, in accordance with previous measurements^40^ (Figure 5f, **Supplementary Figure S9**). Notably mobile β1-ArrayG_16x_ molecules had a 1.9x smaller diffusion coefficient compared to β1-eGFP_2x_. Considering the distribution of diffusivity for integrin β1 spans over ~5 orders of magnitude, a 1.9x reduction may not be functionally relevant, as demonstrated by the ability of β1-ArrayG_16x_ to restore spreading to wild-type levels in a β1 knockout cell line. Moreover, the shape and peaks of the two diffusivity distributions were nearly identical (Figure 5f).

We also checked whether β1-ArrayG_16x_ could respond effectively to an external stimulus, by measuring response to the addition of Mn^2^+, a potent activator of integrins that enhances integrin immobilization.^40^ Stimulation with 2 mM Mn^2^+ for 1 hour immobilized both #x03B2;1-eGFP_2x_ and #x03B2;1-ArrayG_16x_ to a similar extent (Figure 5g, **Supplementary Figure S9**). In summary, ArrayG_16x_ fusion retained the biological functions of integrin β1 in our controls.

For robust detection of states and accurate measurement of state dwell times and transition probabilities, single molecule trajectories need to be longer than the mean state lifetimes.^43^ 20 Hz imaging of β1-ArrayG_16x_ provided an ideal balance between trajectory duration and temporal resolution. At this frequency, although photo-bleaching ultimately limited trajectory length, β1-ArrayG_16x_ significantly increased track duration and throughput compared to previous integrin tracking reports^40, 44^.

Starting with the *a priori* assumption that integrin β1 dwells in three different diffusive states (diffusive, confined, and immobilized)^40^, we segmented trajectories using a variational Bayes treatment of hidden Markov models for single molecule trajectories (vbSPT).^43^ However, a four-state model improved the model evaluation score (**Supplementary Note 4**), revealing four distinct diffusion states (Figure 6a-b). The 4-state model suggested that integrin β1 frequently visits a short-lived binding state, and may subsequently transition into a more confined and longer-lived super-bound state (Figure 6c, **Supplementary Figure S10, Supplementary Note 4**). Such detailed state dynamics have not been previously measured due to the lack of tools for high-throughput acquisition of long-lived trajectories. The multiple binding states are consistent with the multiple conformational states of integrin heterodimers^45^, and the multi-modal binding lifetimes of integrins under tension.^46^

**Figure 6.**
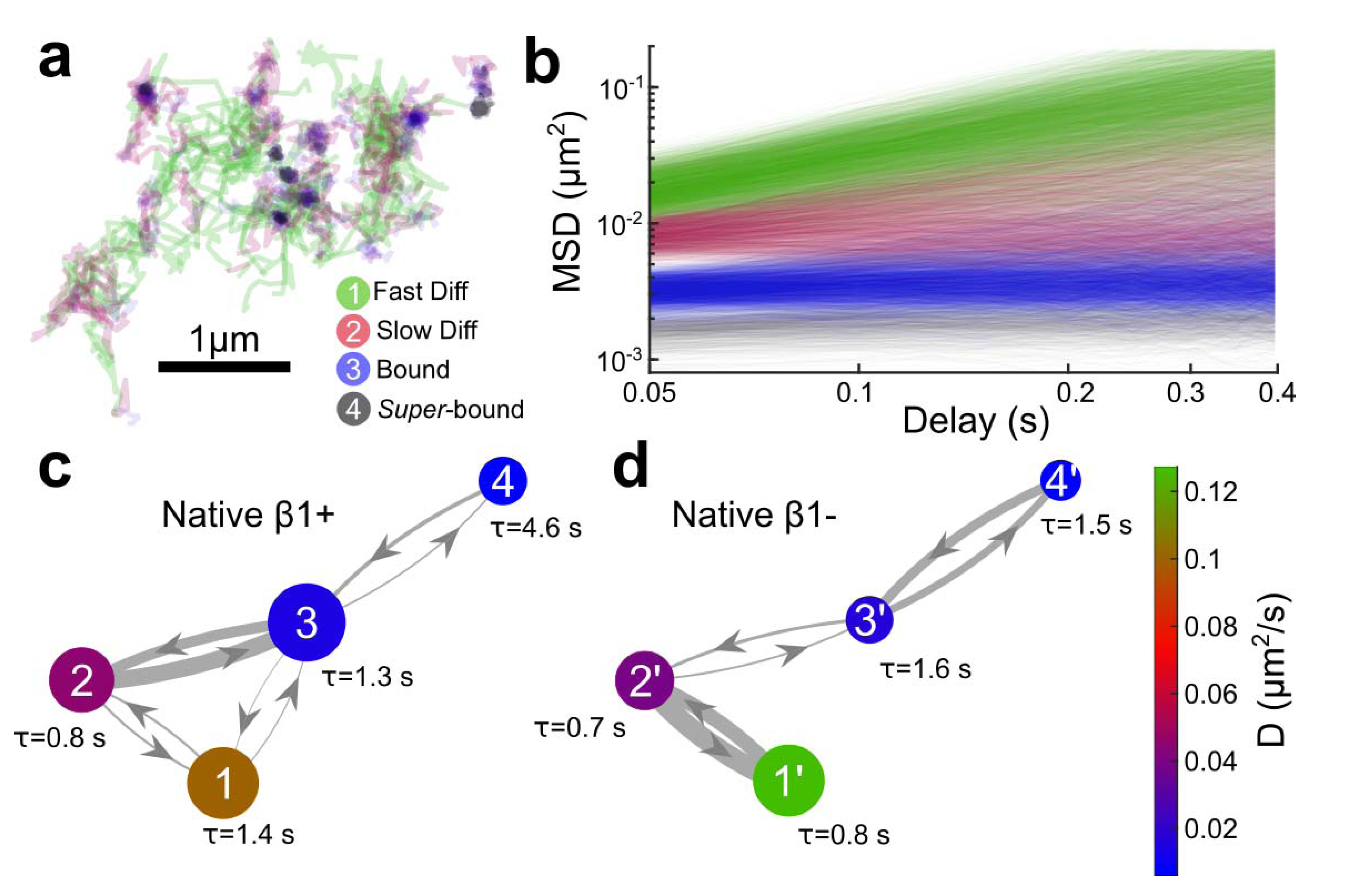
Hidden Markov trajectory segmentation using vbSPT. (**a**) Segmented trajectories color coded by state. (**b**) Overlayed MSD of individual trajectory segments for each state. (**c, d**) Transition state diagrams of β1-ArrayG_16x_ in cells expressing native β1 (pKOαVβ1, N=9,669 trajectories) (**c**) and in β1 null cells (pKOαV, N=6,138 trajectories) (**d**). The area of each state corresponds to the relative occupation (normalized to state 1), the colors correspond to the diffusion coefficient (see colorbar), the width of connecting lines corresponds to the relative transition rate, and the mean state lifetime is written as τ in seconds. Transition rates less than 0.01 per second are ignored.

To test whether the integrin β1 state-landscape depended on local integrin density^47, 48^ we analyzed trajectories of sparsely expressed β1-ArrayG_16x_ in β1-null cells which display impaired contractility (pKOaV β1^-/-^). In contrast to cells expressing native β1, bound state occupancy in β1-null cells was significantly reduced, with more frequent transitions between the fast and slow diffusive states (Figure 6d). These results demonstrate that β1-ArrayG_16x_ dynamics are sensitive to the biological context.

Although vbSPT analysis maybe performed on a large number of short trajectories^43^, the accuracy of the model improved when track lengths were larger than the state lifetimes (**Supplementary note 4**). We found that the various states of integrin mobility had average lifetimes on order of 1 second (ranging from 0.7 to 4.6 seconds), with the super-binder state lasting up to 49 seconds in one instance. This range of lifetimes underscores the importance of long trajectories for distinguishing between states with similar diffusion coefficients, but different lifetimes.

## Outlook

Despite multiple fluorophore recruitment techniques (including SunTag^11^ and Spaghetti-Monster^49^), the field currently lacks optical tagging systems with demonstrably long duration SMT capabilities. The ArrayG system utilizes a nanobody-fluorophore combination that is orthogonal to most eukaryotic complexes. The targets we studied (kinesins, integrins, histones) tolerated the array fusions and we did not see detriments to cell viability, cell division, or intracellular dynamics of the labeled targets. Despite the success with kinesins, integrins, and histones, any new target labeled with the Array tags should be critically evaluated for possible functional impairment.

By exploiting slow exchange of fluorophores at the array, we have found that single molecules of H2B-ArrayGN_24X_ can be optically tracked for very long times, limited only by biological factors such as movement of molecules out of the imaging volume. Note that the traditionally assumed tradeoff between imaging duration and localization precision can be circumvented, via stochastic turnover of FPs on an array, as demonstrated by ArrayG. As shown with the high frequency imaging of kinesins and integrins, the Array tags can also be used to track state transitions at high temporal resolution. The ability to track single biomolecules for temporally unlimited durations may prove useful for addressing questions in assembly, targeting, motility, protein turnover, and cell division. Finally, the fluorogenic ArrayG tag, might allow prolonged SMT in deep tissues where background fluorescence is particularly problematic.

## Materials and Methods

### Array plasmid generation

All mammalian codon optimized Array monomeric unit-DNA were purchased from IDT (integrated DNA technologies) as genes in pUCIDT-Kan plasmid backbone or gBlock gene fragments. All monomeric units had the following structure: *(SalI)-6bp spacer-Scaffold unit-flexible Linker-(XhoI)-6bp spacer-(BamHI).* Using the surrounding SalI-XhoI-BamHI combination, we put the monomer unit through several rounds of repeat building, in which we doubled the number of direct repeats with each cloning cycle, as originally shown by Robinett *et al^8^* After repeat building, the monomer units were moved into a modified Clontech PEGFP-N1 backbone with a Tet-On 3G Tetracycline inducible promoter in place of the CMV promoter (Tet3G backbone). All repeat-containing plasmids were transformed and maintained in MAX Efficiency Stbl2 competent cells (Thermo Fisher Scientific) grown at 30 °C, as long repeats were unstable in other cell types. For generating double stable cell lines, binder-FPs were inserted into the MCS of PiggyBac Cumate switch system (PBQM812A-1, SBI). For the array plasmids, PBQM812A-1 was modified to replace the cumate operator sequence (CuO) with a Tet response element, the EF1-CymR repressor-T2A-Puro cassette was replaced with a custom made EF1-Tet-Activator-T2A-Hygro cassette and the existing MCS was replaced with a custom MCS, such that Arrays could easily be moved from Tet3G backbone plasmids. Cells efficiently transposed for the array and the binder were puromycin and hygromycin positive respectively and were inducible with doxycycline (array) cumate (binder). Doxycycline control was leaky enough to produce arrays in amounts that were ideal for single molecule imaging, when cells were uninduced.

Integrin β1, kinesin and H2B source plasmids were purchased from addgene (Addgene numbers 55064, 15284 and 20972), Vinculin-mCherry was a gift from Johan De Rooij. KIF560 (amino acids 1-560) was generated from full length kinesin. All plasmid sequences are available from NCBI GenBank with the following accession numbers: To be provided by GenBank.

### Cell Culture

HeLa Tet-On® 3G Cell Line (#631183) and U2-OS Tet-On® Cell Line (#631143) were purchased from Clontech. MEF-pKO lines (Shiller *et al.^42^)* were a gift from Alexander Dunn (Dept. of Chemical Engineering, Stanford University). All mammalian cells were cultured on tissue culture plastic in standard tissue culture incubators held at 37°C with 5.0% CO_2_ using DMEM supplemented with 20% fetal bovine serum and penicillin-streptomycin antibiotics. Before plating pKOαV and pKOαVβ1 mouse embryonic fibroblast cell lines, plastic cultureware was incubated with 5μg/mL fibronectin for at least one hour and then washed with PBS before adding culture media and cells.

U2OS cells co-expressing H2B-ArrayGN and mWTGFP were synchronized to G1/S boundary using a double thymidine block procedure. ~30% confluent cells were blocked with 2.5mM Thymidine for ~20 hours followed by a release step lasting 10 hours, followed by a second Thymidine block. Cells were treated with cumate during the 10-hour release step to induce expression of mWTGFP. For imaging cells in prophase, cells were released from the second thymidine block after a period of ~16 hours, incubated for 2 hours in fresh medium, followed by treatment with 10μM RO-33O6 (Sigma) for 10 hours. Release from the RO-33O6 block allowed cells to synchronously enter mitosis.

### Transfections

All transfections were performed using the Neon Transfection system (Thermo Fisher Scientific). HeLa cells were co-transfected with 0.1-0.2 μg of binder plasmids and 4-8 μg of ArrayG or ArrayD plasmids. We generally used two electroporation pulses (pulse voltage 1,005 V, pulse duration 35 ms). pKO-αVμ1 as well as pKO-αV β1^-^/^-^ cells were transfected using the following pulse program: 1 pulse, 1,350 V pulse voltage, 30 ms pulse width. U2OS cells were transfected using 2 pulses of 1,050 V and 30 ms pulse width. Upon transfection, cells were seeded on Mattek dishes coated with fibronectin (Sigma Aldrich) in complete DMEM media without Pen/Strep and with teteracycline free FBS (Thermo Fisher Scientific).

### Cell Line generation

HeLa, U2OS and pKO-aV/αVβ1 cell lines were generated by transfecting cells with the relevant PiggyBAC plasmids and a plasmid expressing transposase following manufacturer’s protocol and then selecting using appropriate antibiotics. For single stable cell line puromycin was the antibiotic of choice where as for double stable cell lines a combination of puromycin and hygromycin was used.

### FACS Analysis

Hela cells stably expressing mwtGFP-P2A-mCherry or mGFP-P2A-mCherry were generated as described above. The cells were analyzed with flow cytometry (BD LSR II) in GFP and mCherry channel along with blank control cells. Only singlets were selected and further analyzed. The ratio of GFP to mCherry per-cell were calculated and displayed after subtracting the background signal from control cells.

### Single GFP quantification for occupancy estimation method 1: confocal calibration

We performed a multi-step calibration of a Zeiss LSM 700 laser scanning confocal so that we could estimate the number of GFPs within each fluorescent spot. First, we measured the bulk fluorescence of an eGFP solution over a wide variety of microscope settings, varying the PMT gain from 450 to 750 volts, confocal pinhole from 41 to 100 μm, and the laser from 1% to 5% power. We fit all the measurements to an arbitrary three-parameter ‘bulk’ model of the form Counts = *A* × laser × (pinhole - B) × gain^C^, allowing us to estimate the solution’s fluorescence at arbitrary instrument settings (**Supplementary Figure S2**).

We measured the bulk per-molar fluorescence of purified eGFP (CellBioLabs), and 0.04 μm green-fluorescent bead standards (5% w/v FluoSpheres, Molecular Probes) by measuring fluorescence at several fluorophore solution concentrations, and several different microscope settings for redundancy (**Supplementary Figure S2f, g**). To measure bead molarity, which is simply particles per unit volume, beads were immobilized by forming a 5% poly-acrylamide gel within a hemacytometer (Incyto) so they could be counted within a known volume (**Supplementary Figure S2e**). eGFP molarity was measured using a UV-Vis spectrophotometer and the published extinction coefficient at 488 nm.^50^ Using these bulk solution calibrations, we can estimate the per-molar fluorescence of both eGFP and the bead solution at arbitrary gain, laser power, and pinhole settings.

Finally, we measured the average fluorescence intensity from single beads adsorbed to glass. Intensity was extracted from each image as the amplitude of a 2D Gaussian fit to each segmented localization. We measured single bead intensity over a range of instrument settings mirroring those used in the bulk calibration, and fit all the measurements to a similar arbitrary four-parameter “single” model of the form Counts = *A* x gain + *B* × laser × (pinhole - C) × D^gain^ (**Supplementary Figure S2c**). This model allowed us to estimate the fluorescence of a single bead at arbitrary instrument settings.

The ratio of fluorescence from a single emitter to a bulk solution of emitters does not depend on the emitter type, Counts_si_ng_le_/Countsb_ul_k = *k,* assuming that emitter concentration and imaging conditions are identical.^51^ Thus, we can solve for the fluorescence intensity of a single eGFP as eGFPsingle = eGFPbulk x (beadsingle/beadbulk). The calibrations allow us to estimate single eGFP intensity at experimental gain, laser power, and pinhole settings, using the bulk per-molar fluorescence from both beads and eGFP calculated at the experimental settings from our ‘bulk’ model, and the fluorescence from single beads at the same settings using our ‘single’ model.

### Single GFP quantification for occupancy estimation method 2: HiLo-TIRFM

In a complementary approach to method 1, we estimated the number of eGFPs within single foci using HiLoTIRFM. As HiLo-TIRFM is far more sensitive than confocal microscopy, we directly measured the fluorescence from single, purified eGFPs (CellBioLabs) adsorbed to plasma-cleaned glass. We added eGFP to a solution of pH 7.4 PBS while providing simultaneous epi-illumination (0.3 kW/cm^2^) 488 nm laser until we began to see blinking events characteristic of single GFPs non-specifically adsorbing to the surface.^44^ The average eGFP intensity was calculated as the amplitude of a 2D Gaussian fit to each discernable spot. We then imaged ArrayG, without changing any instrument settings, and measured the distribution of singlemolecule intensities, again as the amplitude of a 2D Gaussian. We performed a separate eGFP intensity calibration each time each time we quantified ArrayG occupancy.

### TIRF, dual color TIRF and HiLo-TIRFM imaging

Total internal reflection microscopy (TIRF or TIRFM), dual-color TIRFM, and HiLo-TIRFM, were performed on an Olympus CellTIRF system, with independent motor controlled TIRF angles for each fiber-coupled illumination laser, through a 1.49 NA 100× objective and 1.6× Optovar magnifier. Images were recorded on an Andor iXon Plus EMCCD camera at 20 frames per second, unless otherwise noted. Dual-color experiments were imaged through a Photometrics DualView 2 with simultaneous excitation of 488 nm and 561 or 639 nm lasers. All dichroics and filters were purchased from Semrock. All experiments were performed within a heated, CO_2_ controlled incubation chamber set to 37 °C and 5%, respectively. Additional temperature control was provided by a collar-type objective heater, also set to 37 °C.

### Single particle tracking

Single particle tracking (SPT) was performed in two steps, all using custom-written Python and C++ code. First, detectable particles in each frame were identified by performing a Laplacian of Gaussian filter and then thresholding the filtered image based on intensity. A 2D Gaussian centered on each identified particle was then refined onto the raw image data using the Levenberg-Marquardt algorithm, with particle x/y position, amplitude, background height, and in some instances standard deviation as free parameters. All particle detection parameters were validated by eye for each dataset, using a custom GUI. The single particle positions in each frame were then linked into single particle time trajectories using a nearest-neighbor type particle tracking heuristic, with a maximum per-frame jump distance parameter. The algorithm iteratively minimized the jump distance between frames, with no assumed momentum. All single particle trajectories were spot-validated by eye. For signal-to-noise ratio calculations, the signal was defined as the peak height of the Gaussian fit above background level, and noise was defined as the standard deviation of a 5x5 pixel window centered around the particle location after subtracting fitted Gaussian.

### Fractional Brownian Motion

The raw XY,T data were directly fit to a three parameter fBM model (Mathematica version 11.1.1.0, FractionalBrownianMotionProcess[μ, σ, h], ProcessEstimator -> “MaximumLikelihood"]). The first term μ captures constant ‘drift’, σ captures the covariance (= σ^2^ (s^2h^+t^2h^-Abs[t-s]^2h^)/2), and *h* is the Hurst parameter. Note that *h* is simply 1/2 of α, the slope of LogLog transformed MSD. For ease of comparing to the existing literature, we report α rather than the Hurst parameter h.

|  | **0.5 Hz** | **20 Hz** |
| --- | --- | --- |
| Number of cells: | 8 | 11 |
| Datapoints: | 353605 | 103459 |
| Number of trajectories: | 1084 | 445 |
| $\alpha$ Mean: | 0.422115 | 0.434427 |
| SD: | 0.137685 | 0.163548 |
| 95% CI | {0.41391, 0.430321} | {0.41919, 0.449664} |

### Kinesin analysis

Starting with raw SPT data, we first removed stuck particles by filtering out tracks that moved less than 0.4 μm/s as measured by the distance between the first and last frame over the total track length. Freely diffusing particles and tracking failures (common at high particle density) were removed by filtering out tracks that turned too sharply, as measured by the maximum track curvature. Using the filtered subset, kinesin speed and run distance were measured by first smoothing each trajectory with a Savitzky-Golay filter to average away particle localization errors, making a one-dimensional distance axis. Within each trajectory, each localization was then assigned to the closest interpolated point along the smoothed distance axis, giving run distance as a function of time. Each distance versus time trajectory was fit to a simple linear function to measure run speed, and average speed was measured as the peak of the resultant speed distribution. In high time resolution data, free-diffusing and bound kinesins were distinguished by measuring a sliding window average centered around each time point, +/- 10 frames. Regions with a standard deviation larger than 30 nm were considered diffusing, and lower than 30 nm bound. Speed variation along each bound segment was measured as the first derivative of a Savitsky-Golay filter with a window size of 25 frames, and an order of 2. All analysis was performed using custom code in python.

### Cell spreading assay

To achieve a high expression of the integrin β1-ArrayG constructs, cells were induced with cumate at 5x the working concentration diluted from a 10,000x water-soluble stock solution (SBI 300mg/mL QA150A-1) for at least 3 days before assessing cell spreading. The pKOαV + β1-ArrayG_16x_+mGFP cell line was cultured in the absence of tetracycline using Tet-free FBS, to prevent background induction of the Tet-on inducible control of mGFP. Four hours before performing the spreading assay, mGFP expression was induced with Doxycycline at 2 μg/mL. This was done to promote efficient ER processing and subsequent trafficking of integrin to the membrane without GFP binding, and then subsequently assess functionality of the GFP-occupied array on integrin β1 in the membrane. Glass bottomed imaging dishes (No. 1 imaging glass, Mattek) were incubated with 10 μg/mL fibronectin in PBS for 2 hours at 37°C. pKOαV cell lines were then seeded on the fibronectin coated dishes and immediately placed on an confocal microscope (Zeiss LSM 700) with a live imaging incubation chamber equilibrated to 37°C and 6% CO2. Following a 1-hour equilibration period, cells were imaged using the transmission photomultiplier tube (T-PMT) every 1 hour for 5 hours to obtain bright field images of an 8x7 tile (1145 μm x 1000 μm area). Image analysis took place in three steps: (1) stitching of raw image tiles using on-board Zeiss imaging software, (2) local contrast enhancement of stitched images followed by a Sobbel edge finding filter using ImageJ, (3) Segmentation of cells using custom software written in Python to find cell contours. Each cell contour checked by eye to match the raw images, and manually adjusted if needed. Calculation of mean spreading data used data from cells that remained within the image area and did not divide during the experiment. Data presented represents *N =* 50 to 60 cells per condition (specific numbers given in figure) from two independent replicates. The sample sizes used here are similar to previous measurements.^42^

### MSD analysis

Mean-square displacement (MSD) calculations were done using the @msdanalyzer class for MATLAB.^52^ When present, homogenous drift from the microscope stage was corrected using velocity correlations between individual tracks. After drift correction, the MSD was calculated directly from each trajectory.

Diffusion coefficient distribution. Integrins exhibit multiple modes of motion that are apparent in the distribution of diffusion coefficients.^40^ We estimated the diffusion coefficient for individual trajectories, rather than ensemble averaged MSD curves, so that we would not loose information about transitions among molecular behaviors. We calculated the distribution of single molecule diffusion coefficients by fitting Equation 2 over a specified time scale.

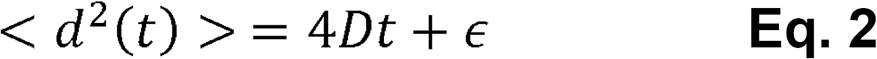

Here, <d^2^> is the MSD of an individual trajectory, *t* is the displacement time scale, *D* is the diffusion coefficient and ε is an offset due to localization error. For all datasets, we used *t* = 0.33 seconds to estimate D. Images of integrin β1-ArrayG_16x_ were recorded at 20Hz, and integrin β1-eGFP_2x_ images required 30Hz recording to accurately capture its dynamics due to the faster diffusion rate. Because the integrin trajectories from ArrayG-labeled molecules sample long time-scales, it is likely that single molecules switch behaviors over a single trajectory, potentially masking the separate behaviors through averaging. Further, immobilized molecules are more easily tracked, and therefore can have much longer trajectory lengths than mobile molecules. To avoid these biases, we divided long trajectories into a series of non-overlapping 1 second segments, and estimated the diffusion coefficient as described above. Smooth distributions were generated using a normal kernel probability density estimation. To compare the relative diffusion rates of eGFP_2x_ and ArrayG_16x_, the log-scaled diffusion coefficient distribution was fit to a two-Gaussian mixture, giving a rate of μm^2^/s for eGFP_2x_ and 0.14 μm^2^/s ArrayG_16x_. Diffusion analysis inside vinculin-mCherry adhesions was performed as follows: (1) registration of GFP and mCherry channels using a rigid-body transformation of the mCherry channel onto the GFP channel from 405 illuminated sample images which produce the same features in each channel,(2)segmentation of time-averaged vinculin-mCherry using a Laplacian of Gaussians filter, and (3)assignment of trajectories that resided within contours of adhesions for 75% of their lifetime as ‘inside’ the adhesion.

### Integrin trajectory segmentation using vbSPT

Integrin β1-ArrayG_16x_ trajectories from either pKO-αV or -αVβ1 cell lines were subjected to segmentation analysis using vbSPT^43^. Settings used were as follows: minimum track length = 20 frames (1 second), initial diffusion coefficient range = [10^-4^, 1] μm^2^/s, initial dwell time range = [0.5, 25] seconds, runs = 16, and default settings for prior distributions of diffusion coefficient, transition rates, and dwell times. Parameter estimations were determined from 100 bootstrap resamplings, besides state dwell times, which were taken as the arithmetic mean based on all trajectories in each state.

### Quantifying the dynamics of binder interaction with cognate arrays using FRAP to determine affinity of each array/binder pair

U2OS tet-on cells were co-transfected with H2B-ArrayGN or H2B-(Snurportin IBB)_4x_-ArrayD and mwtGFP or DHFR-mGFP plasmids respectively. For ensemble FRAP arrays and binders were induced overnight with doxycycline and cumate. All FRAP analysis was carried out on cells with a typical interphase chromatin architecture, using a Zeiss 700 laser scanning confocal microscope enclosed in an incubation chamber set to 5% CO2 and 37 °C. Cells were plated in Mattek glass bottomed dishes coated with fibronectin, and imaged using a 63x oil-immersion objective. Bleaching was carried out using 488 nm laser operating at 100% laser power.

For measuring exchange of mwtGFP on ArrayGN using FRAP, two ROIs per nucleus were generated (~8 μm^2^ per ROI). Background and bleach correction were done using ROIs outside and inside the nucleus, respectively. Corrected intensity curves were fit to a single exponential decay, as free wtGFP contributes little to the fluorescent signal. For DHFR, we spot bleached a ~2 μm diameter circle to follow recovery. Average intensity as a function of time, I(t), was normalized to the average pre-bleach value. DHFR curves were fit with a double exponential function I(t)=A1+A2xEXP(-k1*t)+A3xEXP(-k2*t) (E1) to account for both the fast diffusive free binder-mGFP fraction and the less mobile fraction bound to the cognate scaffold. In case of the control cells expressing mGFP-NLS but no array, the intensity profile was fit with a single exponential function I(t) = A1+A2xEXP(-k1*t) (E2). The fitted parameters were averaged over ~10 repeats to compute means and standard deviations. Half maximum recovery was computed as t_1/2_ = log(0.5)/-k.

### Single molecule FRAP on H2B-ArrayGN

Exchange on arrays was also measured by imaging sparsely expressed H2B-ArrayGN, and performing FRAP on individual localizations. The initial spot intensity was measured by finding the maximum of the localization intensity for each spot in a z-stack of images. Subsequent bleaching was performed under constant illumination and recording 20 Hz images. Intensity measurements were taken at 0, 30, and 60 min post-bleaching using the same procedure as initial spot measurements.

### Data and code availability

All datasets and analysis software are available upon request.

## Acknowledgments

This work was partially supported by the National Institutes of Health (NIH) National Institute Of General Medical Sciences (NIGMS)/National Cancer Institute (NCI) Grant GM77856, NCI Physical Sciences Oncology Center Grant U54CA143836, National Science Foundation Graduate Fellowship Program #DGE-114747, and National Institute Of Biomedical Imaging And Bioengineering (NIBIB)/4D Nucleome Roadmap Initiative 1U01EB021237.

## Author information

### Contributions

R.P.G., J.M.F, W.D., Q.S., and J.T.L. designed research. R.P.G., J.M.F, W.D., and Q.S performed research. J.M.F., W.D., Q.S., and R.P.G. analyzed data. J.T.L., R.P.G., J.M.F, W.D. and Q.S wrote the paper.

### Corresponding author

Correspondence to: Jan Liphardt

